# Reconstruction of a catalogue of genome-scale metabolic models with enzymatic constraints using GECKO 2.0

**DOI:** 10.1101/2021.03.05.433259

**Authors:** Iván Domenzain, Benjamín Sánchez, Mihail Anton, Eduard J. Kerkhoven, Aarón Millán-Oropeza, Céline Henry, Verena Siewers, John P. Morrissey, Nikolaus Sonnenschein, Jens Nielsen

## Abstract

Genome-scale metabolic models (GEMs) have been widely used for quantitative exploration of the relation between genotype and phenotype. Streamlined integration of enzyme constraints and proteomics data into GEMs was first enabled by the GECKO method, allowing the study of phenotypes constrained by protein limitations. Here, we upgraded the GECKO toolbox in order to enhance models with enzyme and proteomics constraints for any organism with an available GEM reconstruction. With this, enzyme-constrained models (ecModels) for the budding yeasts *Saccharomyces cerevisiae, Yarrowia lipolytica* and *Kluyveromyces marxianus* were generated, aiming to study their long-term adaptation to several stress factors by incorporation of proteomics data. Predictions revealed that upregulation and high saturation of enzymes in amino acid metabolism were found to be common across organisms and conditions, suggesting the relevance of metabolic robustness in contrast to optimal protein utilization as a cellular objective for microbial growth under stress and nutrient-limited conditions. The functionality of GECKO was further developed by the implementation of an automated framework for continuous and version-controlled update of ecModels, which was validated by producing additional high-quality ecModels for *Escherichia coli* and *Homo sapiens.* These efforts aim to facilitate the utilization of ecModels in basic science, metabolic engineering and synthetic biology purposes.

## Introduction

Genome-scale metabolic models (GEMs) have become an established tool for systematic analyses of metabolism for a wide variety of organisms^1–6^. Their myriads of applications span from model-driven development of efficient cell factories^3,7–9^, to their utilization for understanding mechanisms underlying complex human diseases^10–12^. One of the most common simulation techniques for enabling phenotype predictions with these models is flux balance analysis (FBA), which assumes that there is balancing of fluxes around each metabolite in the metabolic network. This means that fluxes are constrained by stoichiometries of the biochemical reactions in the network, and that cells have evolved in order to operate their metabolism according to optimality principles^13,14^. Quantitative determination of biologically meaningful flux distribution profiles is a major challenge for constraint-based methods, as optimal phenotypes can be attained by alternate flux distribution profiles^15^, caused by the presence of network redundancies that provide organisms with robustness to environmental and genetic perturbations. This limitation is often addressed by incorporation of experimental measurements of exchange fluxes (secretion of byproducts and uptake of substrates) as numerical flux constraints for the FBA problem. However, such measurements are not readily available for a wide variety of conditions and organisms.

In order to overcome these limitations, the concept of enzymatic limitations on metabolic reactions has been explored and incorporated by several constraint-based methods. Some of these have modelled enzyme demands of metabolic reactions by constraining metabolic networks with kinetic parameters and physiological limitations of cells, such as a crowded intracellular volume^16–18^, a finite membrane surface area for expression of transporter proteins^19^ and a bounded total protein mass available for metabolic enzymes^20–25^. All of these modelling frameworks have been successful at expanding the range of predictions of classical FBA, providing explanations for overflow metabolism and cellular growth on diverse environments for *Escherichia coli*^16–19,21,23,25^, *Saccharomyces cerevisiae*^22,25,26^, *Lactococus lactis*^27^ and even human cells^20,24^. However, these modelling approaches were applied to metabolic networks of extensively studied model organisms, which are usually well represented in specialized resources for kinetic parameters such as the BRENDA^28^ and SABIO RK^29^ databases. Furthermore, collecting the necessary parameters for the aforementioned models was mostly done manually; therefore, no generalized model parameterization procedure was provided as an integral part of these methods.

Enzyme limitations have also been introduced into models of metabolism by other formalisms, for instance, Metabolic and gene Expression models (ME-models), implemented on reconstructions for *E. coli*^30–33^, *Thermotoga maritima*^34^ and *Lactococus lactis*^35^; and resource balance analysis models (RBA), on reconstructions for *E. coil*^36^ and *Bacillus subtilis*^36,37^. These formalisms succeeded at merging genome-scale metabolic networks together with comprehensive representations of macromolecular expression processes, enabling detailed exploration of the constraints that govern cellular growth on diverse environments. Despite the great advances for understanding cell physiology, provided by these modelling formalisms, accuracy on phenotype predictions is compromised by the large number of parameters that are required (rate constants for transcriptional, translational, protein folding and degradation processes), with most of these not being readily available in the literature. Moreover, these models encompass processes that differ radically in their temporal scales (e.g., protein synthesis vs. metabolic rates) and their mathematical representation (presence of non-linear expressions in ME-models), requiring the implementation of more elaborate techniques for numerical simulation.

GECKO, a method for enhancement of GEMs with Enzymatic Constraints using Kinetic and Omics data, was developed in 2017 and applied to the consensus GEM for *S. cerevisiae,* Yeast7^38^. This method extends the classical FBA approach by incorporating a detailed description of the enzyme demands for the metabolic reactions in a network, accounting for all types of enzyme-reaction relations, including isoenzymes, promiscuous enzymes and enzymatic complexes. Moreover, GECKO enables direct integration of proteomics abundance data, if available, as constraints for individual protein demands, represented as enzyme usage pseudo-reactions, whilst all of the unmeasured enzymes in the network are constrained by a pool of remaining protein mass. Additionally, this method incorporates a hierarchical and automated procedure for retrieval of kinetic parameters from the BRENDA database, which yielded a high coverage of kinetic constraints for the *S. cerevisiae* network. The resulting enzyme-constrained model, ecYeast7, was used for successful prediction of the Crabtree effect in wild-type and mutant strains of *S. cerevisiae* and cellular growth on diverse environments and genetic backgrounds, but also provided a simple framework for prediction of protein allocation profiles and study of proteomics data in a metabolic context. Furthermore, the model formed the basis for modeling yeast growth at different temperatures^39^.

Since the first implementation of the GECKO method^38^, its principles of enzyme constraints have been incorporated into GEMs for *B. subtilis*^40^, *E. coli*^41^, *B. coagulant*^42^, *Streptomyces coelicolor*^43^ and even for diverse human cancer cell-lines^2^, showing the applicability of the method even for non-model organisms. Despite the rapid adoption of the method by the constraint-based modelling community, there is still a need for automating the model generation and enabling identification of kinetic parameters for less studied organisms. Here we wanted to build GECKO models for several organisms, and we therefore updated the GECKO toolbox to its 2.0 version. Among other improvements, we generalized its structure to facilitate its applicability to a wide variety of GEMs, and we improved its parameterization procedure to ensure high coverage of kinetic constraints, even for poorly studied organisms. Additionally, we incorporated simulation utility functions, and developed an automated virtual pipeline for update of enzyme-constrained models (ecModels), named ecModels container. This container is directly connected to the original sources of version-controlled GEMs and the GECKO toolbox, offering a continuously updated catalogue of diverse ecModels.

## Results

### Community development of GECKO

To ensure wide application and enable future development by the research community, we established the GECKO toolbox as open-source software, mostly encoded in MATLAB. It integrates modules for enhancement of GEMs with kinetic and proteomics constraints, automated retrieval of kinetic parameters from the BRENDA database (python module), as well as simulation utilities and export of ecModel files compatible with both the COBRA toolbox^44^ and the COBRApy package^45^. The development of GECKO has been continuously tracked in a public repository (https://github.com/SysBioChalmers/GECKO) since 2017, providing a platform for open and collaborative development. The generation of output model files in .txt and SBML L3V1 FBC2^46^ formats enabled the utilization of ecYeastGEM^1^ structure as a standard test to track the effects of any modifications in the toolbox algorithm through the use of the Git version control system, contributing to reproducibility of results and backwards compatibility of code.

Interaction with users of the GECKO toolbox and the ecYeastGEM model has also been facilitated through the use of the GECKO repository, allowing users to raise issues related with the programming of the toolbox or even about conceptual assumptions of the method, which has guided cumulative enhancements. Additionally, technical support for installation and utilization of the toolbox and ecYeastGEM is now provided through an open community chat room (available at: https://gitter.im/SysBioChalmers/GECKO), reinforcing transparent and continuous communication between users and developers.

### New additions to the GECKO toolbox

The first implementation of the GECKO method significantly improved phenotype predictions for *S. cerevisiae’s* metabolism under a wide variety of genetic and environmental perturbations^38^. However, its development underscored some issues, in particular that quantitative prediction of the critical dilution rate and exchange fluxes at fermentative conditions are highly sensitive to the distribution of incorporated kinetic parameters. Although *S. cerevisiae* is one of the most studied eukaryal organisms, not all reactions included in its model have been kinetically characterized. Therefore, a large number of *k*_cat_ numbers measured for other organisms (48.35%), or even non-specific to their reaction mechanism (56.03% of *k*_cat_ values found by introduction of wild cards into E.C. numbers) were needed to be incorporated, in order to fill the gaps in the available data for the reconstruction of the first *S. cerevisiae* ecModel, ecYeast7. Moreover, detailed manual curation of *k*_cat_ numbers was needed for several key enzymes in order to achieve biologically meaningful predictions.

As the BRENDA database^47^ is the main source of kinetic parameters for GECKO, all of the available *k*_cat_ and specific activity entries for non-mutant enzymes were retrieved. In total, 38,280 entries for 4,130 unique E.C. numbers were obtained and classified according to biochemical mechanisms, phylogeny of host organisms and metabolic context (**Supp. file 1**), in order to assess significant differences in distributions of kinetic parameters. This analysis showed that not all organisms have been equally studied. Whilst entries for *H. sapiens, E. coli, R. norvegicus* and *S. cerevisiae* account for 24.02% of the total, very few kinetic parameters are available for most of the thousands of organisms present in the database, showing a median of 2 entries per organism (**Fig. 1A**, **Supp. file 1**). The analysis also showed that kinetic activity can differ drastically, spanning several orders of magnitude even for families of enzymes with closely related biochemical mechanisms (**Fig. 1B**). Finally, it was also observed that *k*_cat_ distributions for enzymes in the central carbon and energy metabolism differ significantly from those in other metabolic contexts across phylogenetic groups of host organisms (life kingdoms, according to the KEGG phylogenetic tree^48^), even without filtering the dataset for entries reported exclusively for natural substrates, as previously done by other studies^49^ (**Fig. 1C**).

**Figure 1.**
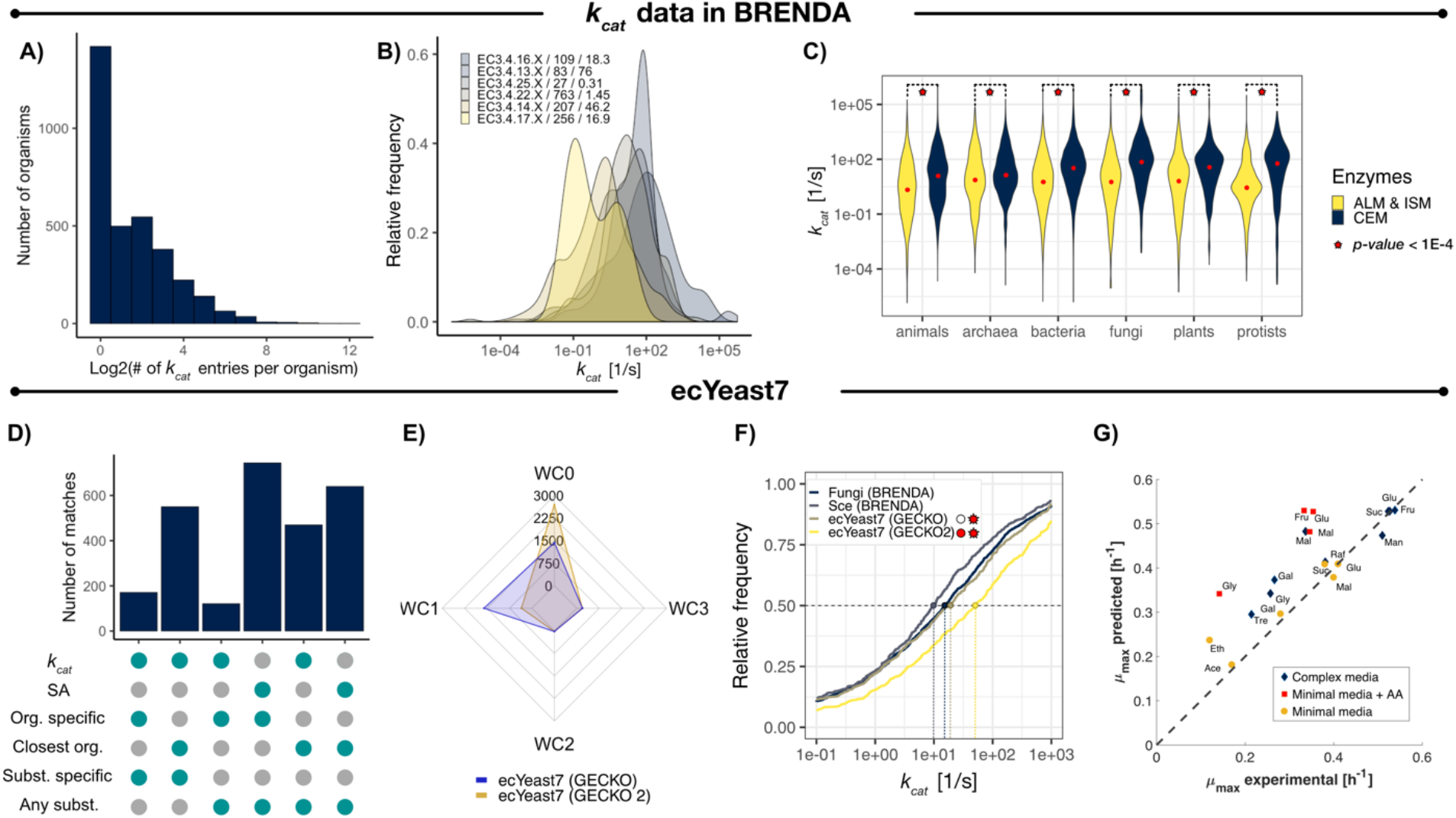
*k_cat_* distributions in BRENDA and ecYeast7. **A)** Number of *k_cat_* entries in BRENDA per organism. **B)** *k_cat_* distributions for closely related enzyme families. Sample size and median values (in s^−1^) are shown after each family identifier. **C)** *k_cat_* distributions for enzymes in BRENDA by metabolic context and life kingdoms. Median values are indicated by red dots in each distribution, statistical significance (under a Kolmogorov-Smirnov test) is indicated by red stars for each pair of distributions for a given kingdom. CEM – central carbon and energy metabolism; ALM – Amino acid and lipid metabolism; ISM – Intermediate and secondary metabolism. **D)** Number of *k*_cat_ matches in ecYeast7 per assignment category (GECKO 2.0). **E)** Comparison of the number of *k_cat_* matches for E.C. numbers with 0, 1, 2 and 3 introduced wild-cards by GECKO 2.0 and GECKO *k_cat_* matching algorithms. **F)** Cumulative *k_cat_* distributions for: all *S. cerevisiae* entries in BRENDA, all entries for fungi in BRENDA, ecYeast7 enhanced by GECKO and ecYeast7 enhanced by GECKO 2.0. Colored points and vertical dashed lines indicate the value for the median value for each distribution. Statistical significance under a Kolmogorov-Smirnov test of the matched *k_cat_* distributions when compared to all entries for *S. cerevisiae* and fungi, is shown with red circles and stars, respectively. *p-values* below 1×10^−2^ are indicated with red. **G)** Prediction of batch maximum growth rates on diverse media with ecYeast7 enhanced by GECKO 2.0. Glu – glucose, Fru – fructose, Suc – sucrose, Raf – raffinose, Mal – maltose, Gal – galactose, Tre – trehalose, Gly – glycerol, Ace – acetate, Eth – ethanol.

As *k*_cat_ numbers depend on biochemical mechanisms, metabolic context and phylogeny of host organisms, a modified set of hierarchical *k*_cat_ matching criteria was implemented as part of GECKO 2.0. The modified parameterization procedure enables the incorporation of kinetic parameters that have been reported as *specific activities* in BRENDA when no *k*_cat_ is found for a given query (as the specific activity of an enzyme is defined as its *k*_cat_ over its molecular weight), adding 8,118 new entries to the catalogue of kinetic parameters in the toolbox. A phylogenetic distance-based criterion, based on the phylogenetic tree available in the KEGG database^48^, was introduced for cases in which no organism-specific entries are available for a given query in the kinetic parameters dataset. A comparison of the new *k*_cat_ matching criteria with their predecessor set is shown in **Supp. file 2**.

In order to assess the impact of the modified *k*_cat_ assignment algorithm on an ecModel, ecYeast7 was reconstructed using both the first and the new version of the GECKO toolbox (GECKO 2.0). A classification of the matched *k*_cat_ numbers according to the different levels of the new matching algorithm is provided in **Fig. 1D**. The incorporation of specific activity values in the parameter catalogue increased the number of kinetic parameters matched to complete E.C. numbers (no added wild cards) from 1432 to 2696 (**Fig. 1E**). Moreover, the implementation of the phylogenetic distance-based criterion yielded a distribution of kinetic parameters that showed no significant differences when compared to the values reported in BRENDA for all fungi species, in contrast to the kinetic profile matched by the previous algorithm (*p-values* <10^−10^ and <10^−7^, when compared to the BRENDA fungi and *S. cerevisiae* distributions, respectively, under a Kolmogorov-Smirnov test) (**Fig. 1F**). The quality of phenotype predictions for the ecYeast7 model enhanced by GECKO2.0 was evaluated by simulation of batch growth in 19 different environments, with an average relative error of 29.22% when compared to experimental data (**Fig. 1G**).

The introduction of manually curated *k*_cat_ numbers in a metabolic network has been proven to increase the quality of phenotype predictions for *S. cerevisiae*^22,25,38^; nevertheless, this is an intensive and time consuming procedure that is hard to ensure for a large number of models subject to continuous modifications. In order to ensure applicability of the GECKO method to any standard GEM, a unified procedure for curation of kinetic parameters was developed based on parameter sensitivity analysis. For automatically generated ecModels that are not able to reach the provided experimental value for maximum batch growth rate, an automatic module performs a series of steps in which the top enzymatic limitation on growth rate is identified through the quantification of enzyme control coefficients. For such enzymes, the E.C. number is obtained and then its correspondent *k*_cat_ value is substituted by the highest one available in BRENDA for the given enzyme class. This procedure iterates until the specific growth rate predicted by the model reaches the provided experimental value.

Finally, as the first version of the toolbox relied on the structure and nomenclature of the model Yeast7, its applicability to other reconstructions was not possible in a straightforward way. In order to provide compatibility with any other GEM, based on COBRA^44^ or RAVEN^50^ formats, all of the organism-specific parameters required by the method (experimental growth rate, total protein content, organism name, names and identifiers for some key reactions, etc.) can be provided in a single MATLAB initialization script, minimizing the modifications needed for the generation of a new ecModel.

### ecModels container: an automatically updated repository

Several GEMs that have been published are still subject to continuous development and maintenance^1–3,5,6^, this renders GEMs to be dynamic structures that can change rapidly. In order to integrate such continuous updates into the enzyme constrained version of a model in an organized way, an automated pipeline named ***ecModels container*** was developed.

The ecModels container is a continuous integration implementation whose main functionality is to provide a catalogue of ecModels for several relevant organisms that are automatically updated every time a modification is detected either in the original GEM source repository or in the GECKO toolbox, i.e. new releases in their respective repositories. The pipeline generates ecModels in different formats, including the standard SBML and MATLAB files, and stores them in a container repository (https://github.com/SysBioChalmers/ecModels) in a version controlled way, requiring minimal human interaction and maintenance. The GECKO toolbox ensures the creation of functional and calibrated ecModels that are compatible with the provided experimental data (maximum batch growth rate, total protein content of cells and exchange fluxes at different dilution rates as an optional input). This whole computational pipeline is illustrated in **Fig. 2**. Further description of the ecModels container pipeline functioning is included in in the **Materials and Methods** section.

**Figure 2.**
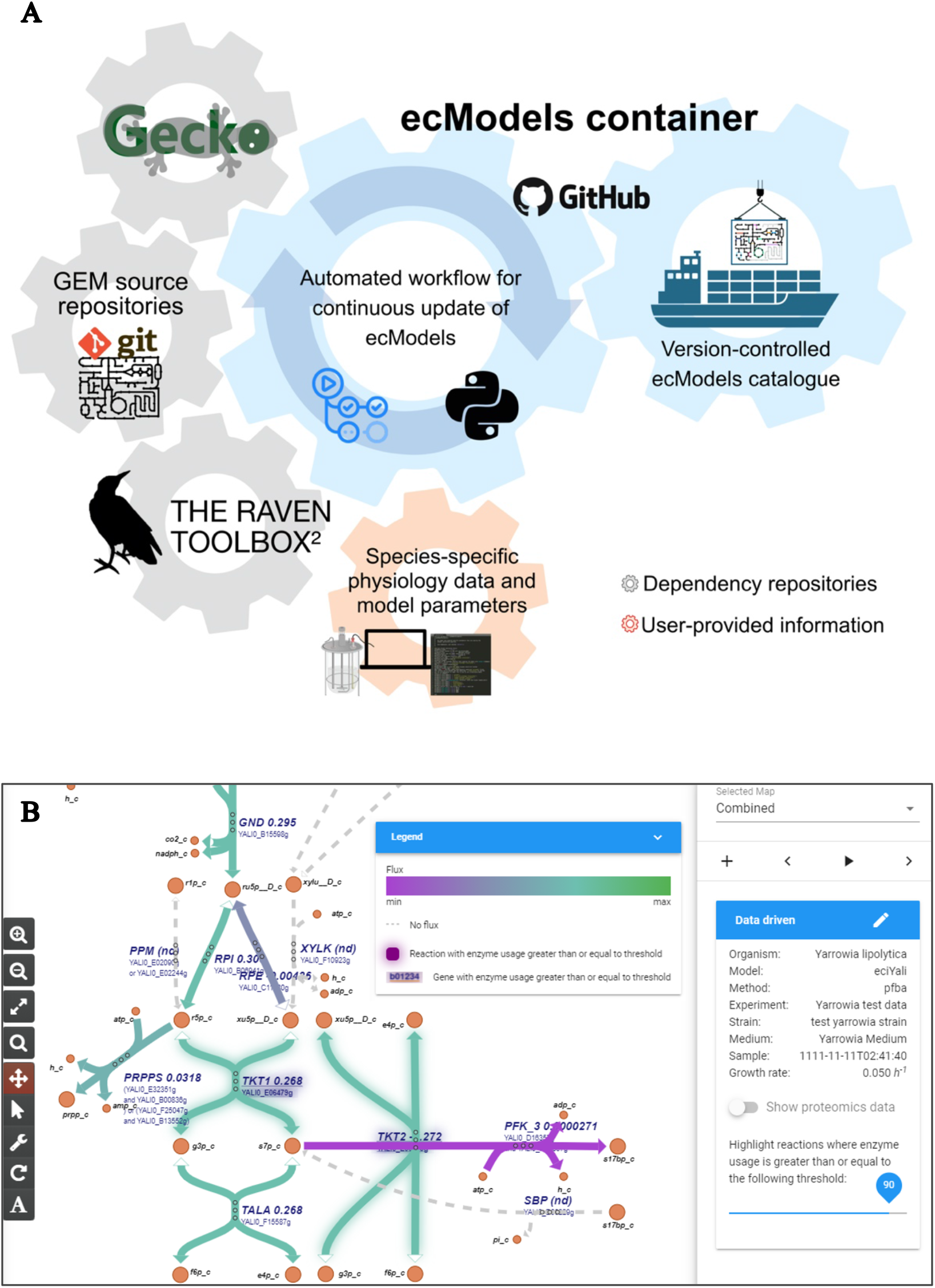
Extending utilization of ecModels. **A)** ecModels container: Integrated pipeline for continuous and automated update of ecModels. **B)** Implementation of GECKO simulations in the Caffeine platform (https://caffeine.dd-decaf.eu/) for visualization of enzyme usage. The color of the arrows corresponds to the value of the corresponding fluxes. Genes or reactions connected to enzymes with a usage above 90% are highlighted with a glow around the corresponding text or arrow, respectively. The chosen usage threshold to highlight can be tuned with the slider on the right.

### A catalogue of new ecModels

Following the aforementioned additions to the GECKO toolbox, that have allowed its generalization, we used the toolbox for the reconstruction of four new ecModels from previously existing high-quality metabolic network reconstructions: *i*Yali4, for the oleaginous yeast *Yarrowia lipolytica*^5^; *i*SM996, for the thermotolerant yeast *Kluyveromyces marxianus*^6^; *i*ML1515, for the widely studied bacterium *E. colŕ;* and Human1, being the latest and largest network reconstruction available for studying *H. sapiens* metabolism^2^. For the microbial models, all model parameters were calibrated according to the provided experimental data, generated by independent studies^4,51–53^, yielding functional ecModels ready for simulations. Size metrics for these models can be seen in **Table 1**.

**Table 1.**
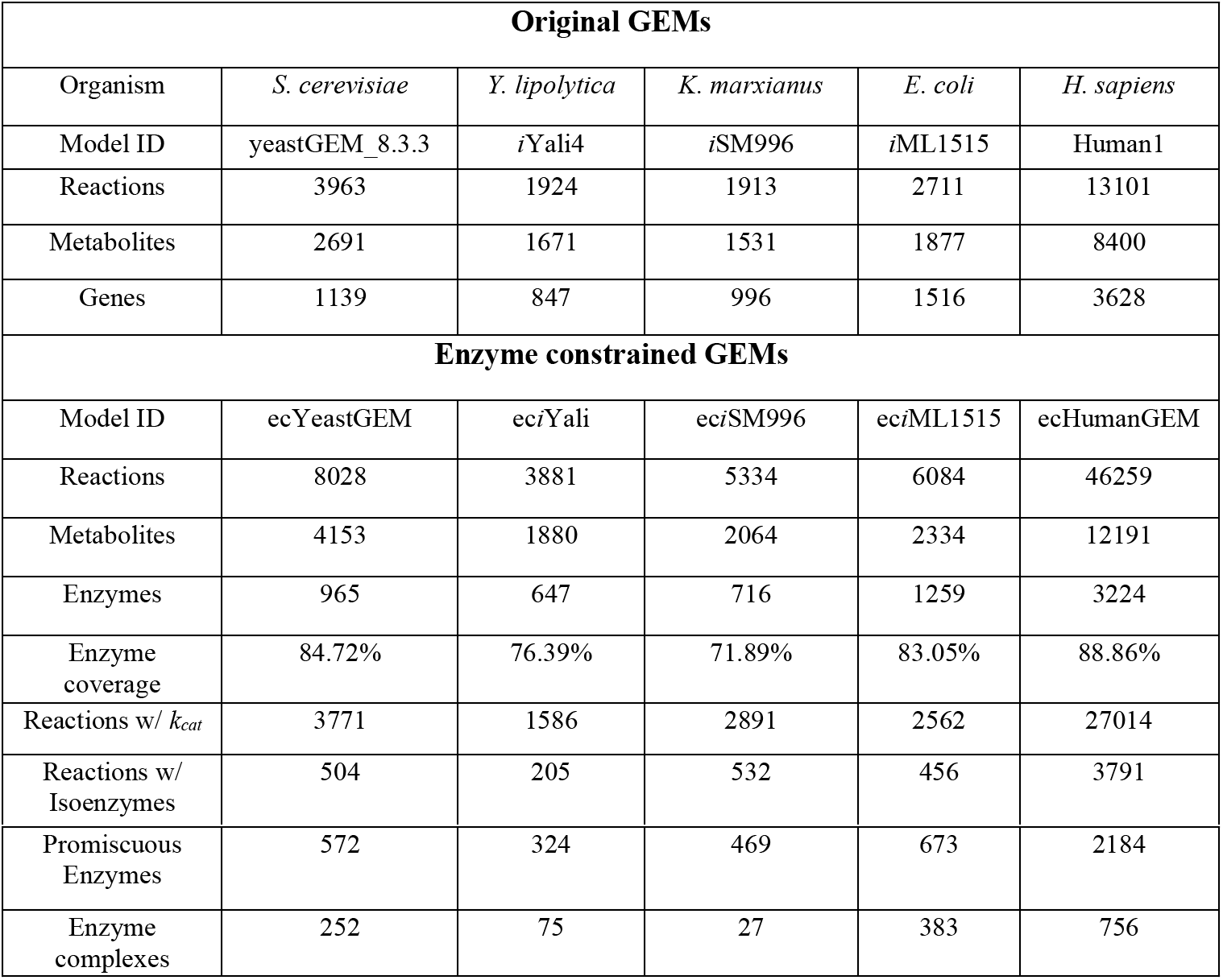
Size metrics summary for the ecModels catalogue.

These ecModels, together with ecYeastGEM, are hosted in the ecModels container repository for their continuous and automated update every time that a version change is detected either in the original model source or in the GECKO repository. In the case of microbial species, two different model structures are provided: ***ecModel***, which has unbounded individual enzyme usage reactions ready for incorporation of proteomics data; and ***ecModel_batch*** in which all enzyme usage reactions are connected to a shared protein pool. This pool is then constrained by experimental values of total protein content, and calibrated for batch simulations using experimental measurements of maximum batch growth rates on minimal glucose media, thus providing a functional ecModel structure ready for simulations.

For ecHumanGEM just the unbounded ecModel files are provided, as this is a general network of human metabolism, containing all reactions from any kind of human tissue or cell type for which evidence is available, and therefore not suitable for numerical simulation. As *H. sapiens* is the most represented organism in the BRENDA database, accounting for 11% of the total number of available *k*_cat_ values (**Supp. file 1**), kinetic parameters from other organisms were not taken into account for its enhancement with enzyme constraints. ecHuman1 provides the research community with an extensive knowledge base that represents a complete and direct link between genes, proteins, kinetic parameters, reactions and metabolites for human cells in a single model structure, subject to automated continuous update by the ***ecModels container*** pipeline.

### Visualization of GECKO simulations in the Caffeine platform

We implemented simulations with ecModels in Caffeine, an open-source software platform for cell factory design. Caffeine, publicly available at http://caffeine.dd-decaf.eu, allows user-friendly simulation and visualization of flux predictions made by genome-scale metabolic models. Several standard modelling methods are already included in the platform, such as ^13^C fluxomics data integration, and simulation of gene deletion and/or overexpression, to interactively explore strain engineering strategies. In order to allow for GECKO simulations, we added a new feature to the platform for uploading enzyme-constrained models and absolute proteomics data. Additionally, we added a simulation algorithm that recognizes said models, and overlays the selected proteomics data on them, leaving out data that makes the model unable to grow at a pre-specified growth rate. After these inclusions to the platform, enzyme usage can now be computed on the fly and visualized on metabolic maps (**Fig. 2B**), to identify potential metabolic bottlenecks in a given condition. The original proteomics data can be visualized as well, to identify if the specific bottleneck is due to a lack of enzyme availability, or instead due to an inefficient kinetic property. This will suggest different metabolic engineering strategies to the user: if the problem lies in the intracellular enzyme levels, the user can interpret this as a recommendation for overexpressing the corresponding gene, whereas if the problem lies in the enzyme efficiency, the user could assess introducing a heterologous enzyme as an alternative.

### GECKO simulation utilities

As ecModels are defined in an irreversible format and incorporate additional elements such as enzymes (as new pseudo-metabolites) and their usages (represented as pseudo-reactions), they might sometimes not be directly compatible with all of the functionalities offered by currently available constraint-based simulation software^44,45,50,54,55^. We therefore added several new features to the GECKO toolbox that allow the exploration and exploitation of ecModels. These include utilities for: 1) basic simulation and analysis purposes, 2) accessible retrieval of kinetic parameters, 3) automated generation of condition-dependent ecModels with proteomic abundance constraints, 4) comparative flux variability analysis between a GEM and its ecModel counterpart, and 5) prediction of metabolic engineering targets for enhanced production with an implementation of the FSEOF method^56^ for ecModels. Detailed information about the inputs and outputs for each utility can be found on their respective documentation, available at: https://github.com/SysBioChalmers/GECKO/tree/master/geckomat/utilities. All of these utilities were developed in MATLAB due to their dependency on some RAVEN toolbox functions^50^.

### Predicting microbial proteome allocation in multiple environments

In order to test the quality of the phenotype predictions of an ecModel automatically generated by the ***ecModels container*** pipeline, batch growth under 11 different carbon sources was simulated with ec*i*ML1515 for *E. coli.* **Figure 3A** shows that, for all carbon sources, growth rates were predicted at the same order of magnitude as their corresponding experimental measurements, with the most accurate predictions obtained for growth on D-glucose, mannose and D-glucosamine. Furthermore, batch growth rate and protein allocation predictions, using no exchange flux constraints, were compared between ec*i*ML1515 and the *i*JL1678 ME-model^32^, the latter accounting for both metabolism and macromolecular expression processes. The sum squared error (SSE) for batch growth rate predictions across the 11 carbon sources using ec*i*ML1515 was 0.27, a drastic improvement when compared to the 1.21 SSE of *i*JL1678 ME-model predictions^32^. **Figure 3B** shows the predicted total proteome needed by cells to sustain the provided experimental growth rates for the same 11 environments. Notably ec*i*ML1515 predicts values that lie within the range of predictions of the *i*JL1678 ME-model (from the optimal to the generalist case) for 10 out of the 11 carbon sources (see Materiales and Methods for simulation details). This shows that the new version of the GECKO toolbox ensures the generation of functional ecModels that can be readily used for simulation of metabolism, due to its systematic parameter flexibilization step which reduces the need of extensive manual curation for new ecModels. Furthermore, *i*ML1515 is a model available as a static file at the BiGG models repository^57^; therefore, its integration to the ecModels container for continuous update demonstrates the flexibility of our pipeline, regarding compatibility with original GEM sources, which can be provided as a link to their *git-*based repositories or even as static URLs.

**Figure 3.**
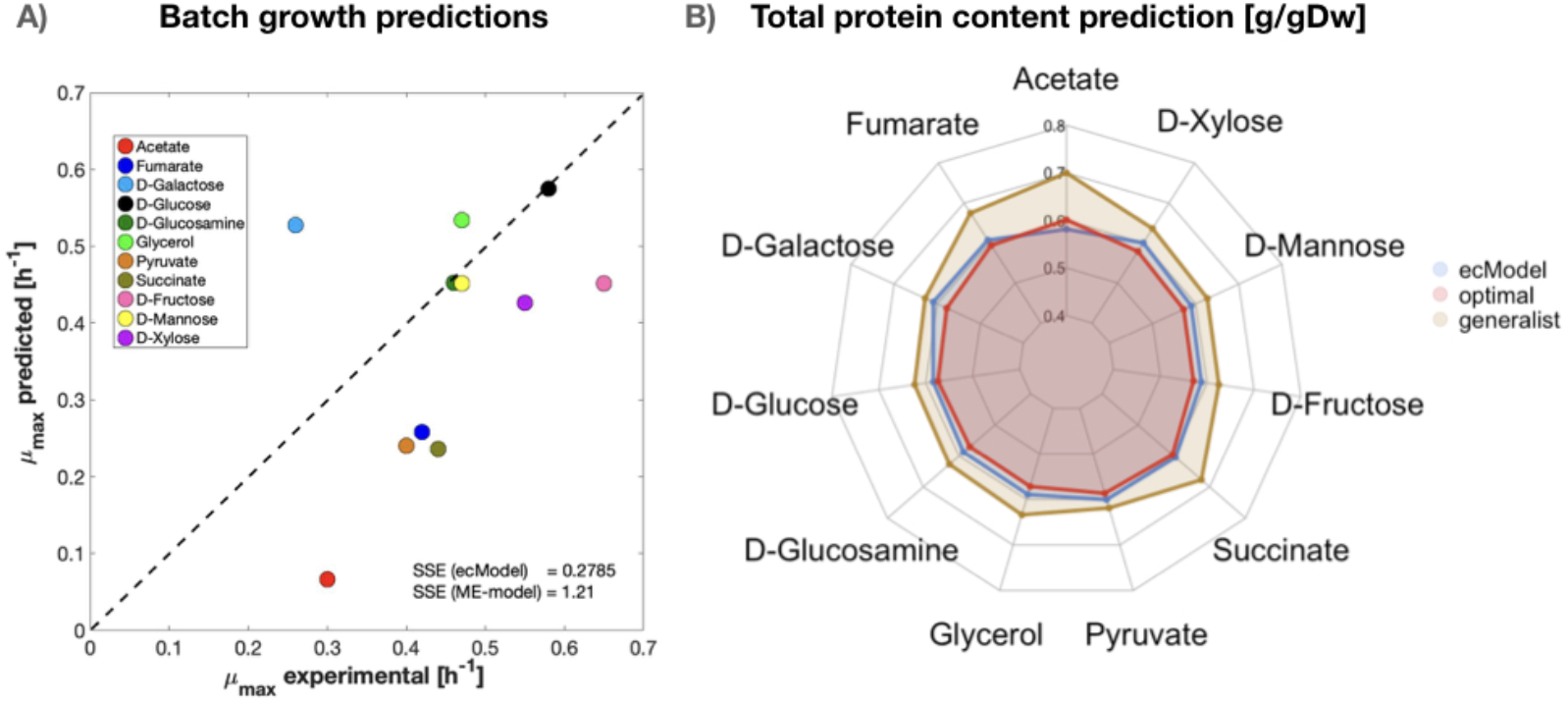
Comparison of predictive capabilities between ec*i*ML1515 and ME-*i*JL1678 for *E. coli.* **A)** Maximum batch growth rate predictions on minimal media with diverse carbon sources. Squared-sum errors when compared to experimental values are shown for both ec*i*ML1515 and ME-*i*JL1678. **B)** Prediction of total protein content in the cell by ec*i*ML1515 and ME-*i*JL1678 using the optimal and generalist approaches.

### Proteomics constraints refine phenotype predictions for multiple organisms and conditions

The previously mentioned module for integration of proteomics data generates a condition-dependent ecModel with proteomics constraints for each condition/replicate in a provided dataset of absolute protein abundances [mmol/gDw]. Even though absolute quantification of proteins is becoming more accessible and integrated into systems biology studies^58–62^, a major caveat of using proteomics data as constraints for quantitative models is their intrinsic high biological and technical variability^63^, therefore some of the incorporated data constraints need to be loosened in order to obtain functional ecModels. When needed, additional condition-dependent exchange fluxes of byproducts can also be used as constraints in order to limit the feasible solution space. A detailed description of the proteomics integration algorithm implemented in GECKO is given in **Supp. file 2**.

The new proteomics integration module was tested on the three ecModels for budding yeasts available in ecModels container (ecYeastGEM, ec*i*Yali, ec*i*SM996). We measured absolute protein abundances for *S. cerevisiae, Y. lipolytica* and *K. marxianus,* grown in chemostats at 0.1 h ^-1^ dilution rate and subject to several experimental conditions (high temperature, low pH and osmotic stress with KCl)^64^, and incorporated these data into the ecModels as upper bounds for individual enzyme usage pseudo-reactions. Then, exchange fluxes for CO_2_ and oxygen corresponding to the same chemostat experiments were used as a comparison basis to evaluate quality of phenotype predictions. For each organism-condition pair, 3 models were generated and compared in terms of predictions: a pure stoichiometric metabolic model, an enzyme-constrained model with a limited shared protein pool, and an enzyme-constrained model with proteomics constraints. It was found that the addition of the enzyme pool constraint enables major reduction of the relative error in prediction of gaseous exchange fluxes in some of the studied conditions. Additionally, the incorporation of individual protein abundance constraints improves even further the predictive accuracy of gaseous exchanges, for 10 out of the 11 evaluated cases (**Fig. 4A-C**).

**Figure 4.**
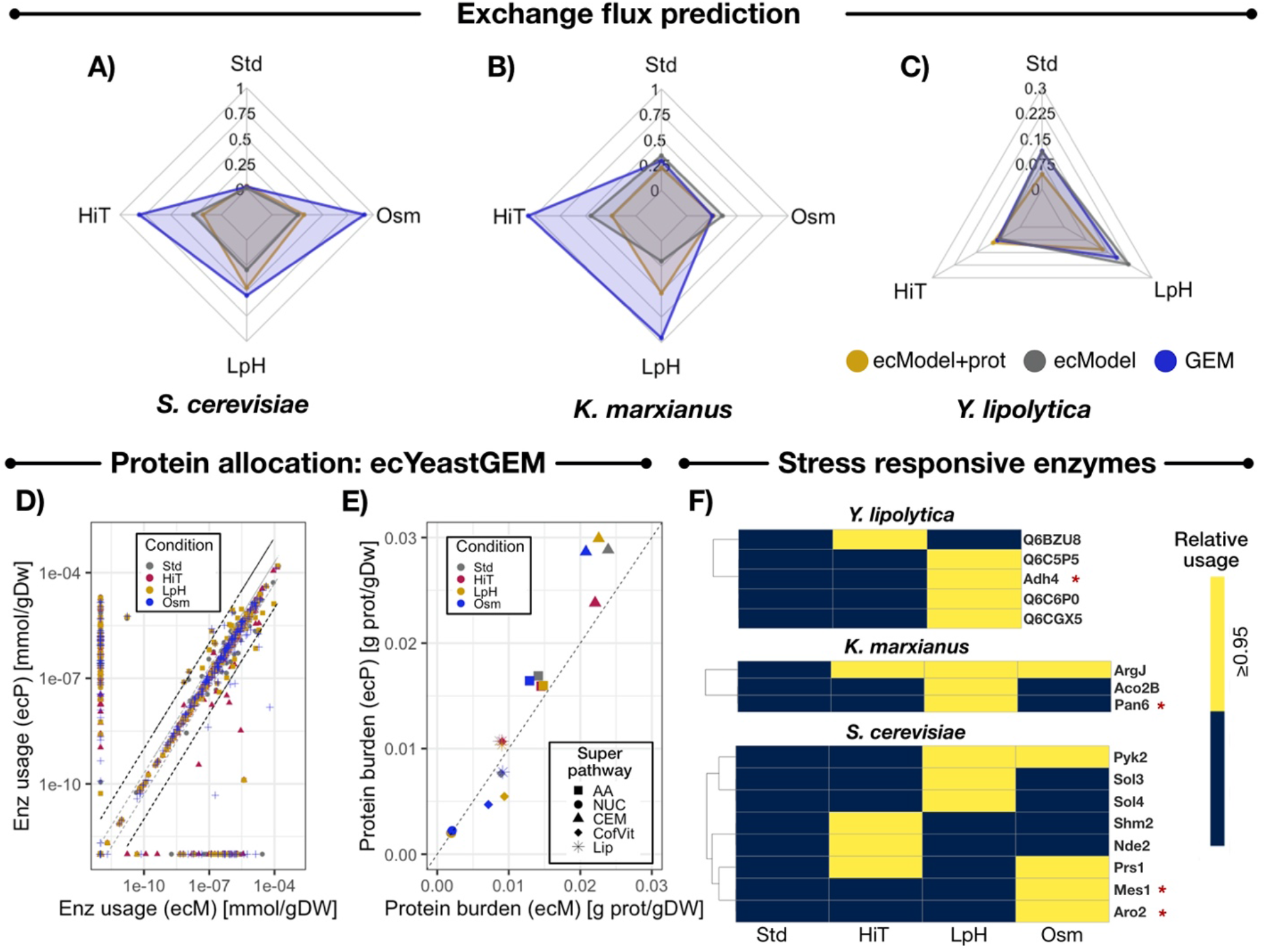
Evaluation of proteomics-constrained ecModels. Comparison of median relative error in prediction of exchange fluxes for O_2_ and CO_2_ by GEMs, ecModels and proteomics-constrained ecModels across diverse conditions (chemostat cultures at 0.1 h^−1^ dilution rate) for **A)** *S. cerevisiae* **B)** *K. marxianus* **C)** *Y. lipolytica.* **D)** Comparison of absolute enzyme usage profiles [mmol/gDw] predicted by ecYeastGEM (ecM) and ecY eastGEM with proteomics constraints (ecP) for several experimental conditions. The region between the two dashed grey lines indicates enzyme usages predicted in the interval 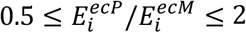, the region between the two dashed black lines indicates enzyme usages predicted in the interval 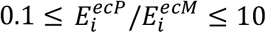 when comparing the two ecModels. **E)** Protein burden for different superpathways predicted by ecYeastGEM (ecM) and ecYeastGEM with proteomics constraints (ecP). **F)** Highly saturated enzymes at different stress conditions for *S. cerevisiae*, *K. marxianus* and *Y. lipolytica* predicted by their corresponding ecModels constrained with proteomics data. Yellow cells indicate condition-responsive enzymes (*relative usage* ≥ 0.95). Red asterisks indicate enzymes conserved as single copy orthologs across the three yeast species. Std – Reference condition, HiT – High temperature condition, LpH – Low pH condition, Osm – Osmotic stress condition, AA – amino acid metabolism, NUC – nucleotide metabolism, CEM – central carbon and energy metabolism, CofVit – cofactor and vitamin metabolism, Lip – lipid and fatty acid metabolism.

The impact of incorporating enzyme and proteomics constraints on intracellular flux predictions was further assessed by mapping all condition-dependent flux distributions from the tested ecModels to their corresponding reactions in the original GEMs. In general, metabolic flux distributions showed high similarity when comparing ecModel to GEM predictions (**Fig. S1**), as 70-90% of the active fluxes were predicted within the interval of 0.5 < fold-change < 2 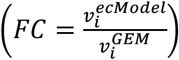 across all conditions (**Fig. S2 A-C**, **Supp. file 3**). In addition, principal component analysis on absolute enzyme usage profiles predicted by ecModels revealed that, at low dilution rates, predictions of enzyme demands are mostly defined by the selected set of imposed constraints (shared protein pool vs. proteomics constraints) rather than by environmental condition, i.e. exchange fluxes (**Fig. S2 D-F)**. However, more straightfroward comparison of the models’ predictions, by pairwise comparison of predicted absolute enzyme usage profiles, showed that 60 – 80% of the predicted enzyme usages lie within a range of 0.5 < fold-change < 2, when comparing ecModels predictions with and without proteomics constraints, across organisms and conditions (**Fig. 4D**, **Fig. S2 G-I** and **Supp. file 3**). It was observed that the incorporation of proteomics constraints induces a drastic differential use for a considerable amount of enzymes, as 12-21% of enzyme usages were predicted as either enabled or disabled by these constraints across all the simulated conditions, showing slight enrichment for enabled alternative isoenzymes for already active reactions (**Supp. file 3**). This suggests that upper bounds on enzyme usages induce differentiated utilization of isoenzymes, reflecting well why isoenzymes have been maintained throughout evolution.

The explicit inclusion of enzymes into GEMs by the GECKO method enables prediction of enzyme demands at the protein, reaction and pathway levels. Total protein burden values predicted by ecModels for several relevant metabolic superpathways (central carbon and energy metabolism, amino acid metabolism, lipid and fatty acid metabolism, cofactor and vitamin metabolism and nucleotide metabolism, according to the KEGG metabolic subsystems^48^), showed that central carbon and energy metabolism is the most affected sector in the ecYeastGEM network by integration of proteomics constraints, as protein burden predictions were higher, at least by 20%, for 3 out the 4 simulated conditions when compared with predictions of the ecYeastGEM without proteomics data (**Fig. 4E**).

Relative enzyme usages, estimated as predicted absolute enzyme usage over enzyme abundance for all of the measured enzymes in an ecModel 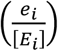, can be understood as the saturation level of enzymes in a given condition. In order to analyze the metabolic mechanisms underlying long-term adaptation to stress in budding yeasts, relative enzyme usage profiles were computed from all the previous simulations of ecModels with proteomics constraints. Enzymes that display fold-changes higher than 1 for both absolute abundance and their saturation level, when comparing predicted usage profiles between stress and reference conditions, suggest regulatory mechanisms on individual proteins that contribute to cell growth on the anlyzed stress condition. **Figure 4F** shows all of the enzymes that were identified as responsive to environmental stress in this study, displaying enrichment for enzymes involved in biosynthesis of diverse amino acids and folate metabolism.

A further mapping of all enzymes in these ecModels to a list of 2,959 single copy protein-coding gene orthologs across the three yeast species^64^ found 310 core proteins across these ecModels. Principal component analysis revealed that variance on absolute enzyme usages and abundance profiles for these core proteins is mostly explained by differences in the metabolic networks of the different species rather than by environmental conditions (**Fig. S3 B-C**), reinforcing previous results suggesting that, despite being phylogenetically related, their long-term stress responses at the molecular level have evolved independently after their divergence in evolutionary history^64^.

#### Exploring the solution space reduction

A major limitation in the use of GEMs is the high variability of flux distributions for a given cellular objective when implementing flux balance analysis, as this requires solving largely underdetermined linear systems through optimization algorithms^15,65^. This limitation has usually been overcome with incorporation of measured exchange fluxes as constraints. However, these data are typically sparse in the literature. Previous studies explored the drastic reduction in flux variability ranges of ecModels for *S. cerevisiae* and 11 human cell-lines when compared to their original GEMs due to the addition of enzyme constraints^1,2,38^. However, the irreversible format of ecModels (forward and backwards reactions are split in order to account for enzyme demands of both directions) hinders their compatibility with the flux variability analysis (FVA) functions already available in COBRA^44^ and RAVEN^50^ toolboxes. As a solution to this, an FVA module was integrated to the utilities repertoire in GECKO, whose applicability has been previously tested on studies with ecModels for *S. cerevisiae*^1^ and human cell lines^2^. This module contains the necessary functions to perform FVA on any set of reactions of an ecModel, enabling also a direct comparison of flux variability ranges between an ecModel and its GEM counterpart in a consistent way (see **Supp. file 2**).

The FVA utility was applied on three different ecModels of microbial metabolism and their correspondent GEMs (*i*ML1515, *i*Yali4 and *i*SM996). In all cases the FVA comparisons were carried out for both chemostat and batch growth conditions in order to span different degrees of constraining of the metabolic networks (0.1 h^−1^ dilution rate and minimal glucose uptake rate fixed for chemostat conditions; biomass production fixed to experimental measurements of *μ_max_* and unconstrained uptake of minimal media components, for batch conditions). Cumulative distributions for flux variability ranges for all explored ecModels and GEMs are shown in **Figure 5**, in which it can be seen that median flux variability ranges are much reduced for all ecModels and conditions, especially at high growth rates where enzyme constraints reduce the variability range 5-6 orders of magnitude when compared to pure GEMs. The cumulative distributions also show a major reduction in the amount of totally variable fluxes (reactions that can carry any flux between −1000 to 1000 mmol/gDw h), which are an indicator of undesirable futile cycles present in the network due to lack of thermodynamic and enzyme cost information^66–68^. For high growth rates, the amount of totally variable fluxes accounts for 3-12% of the active reactions in the analysed GEMs, in contrast to their corresponding ecModels in which such extreme variability ranges are completely absent.

**Figure 5.**
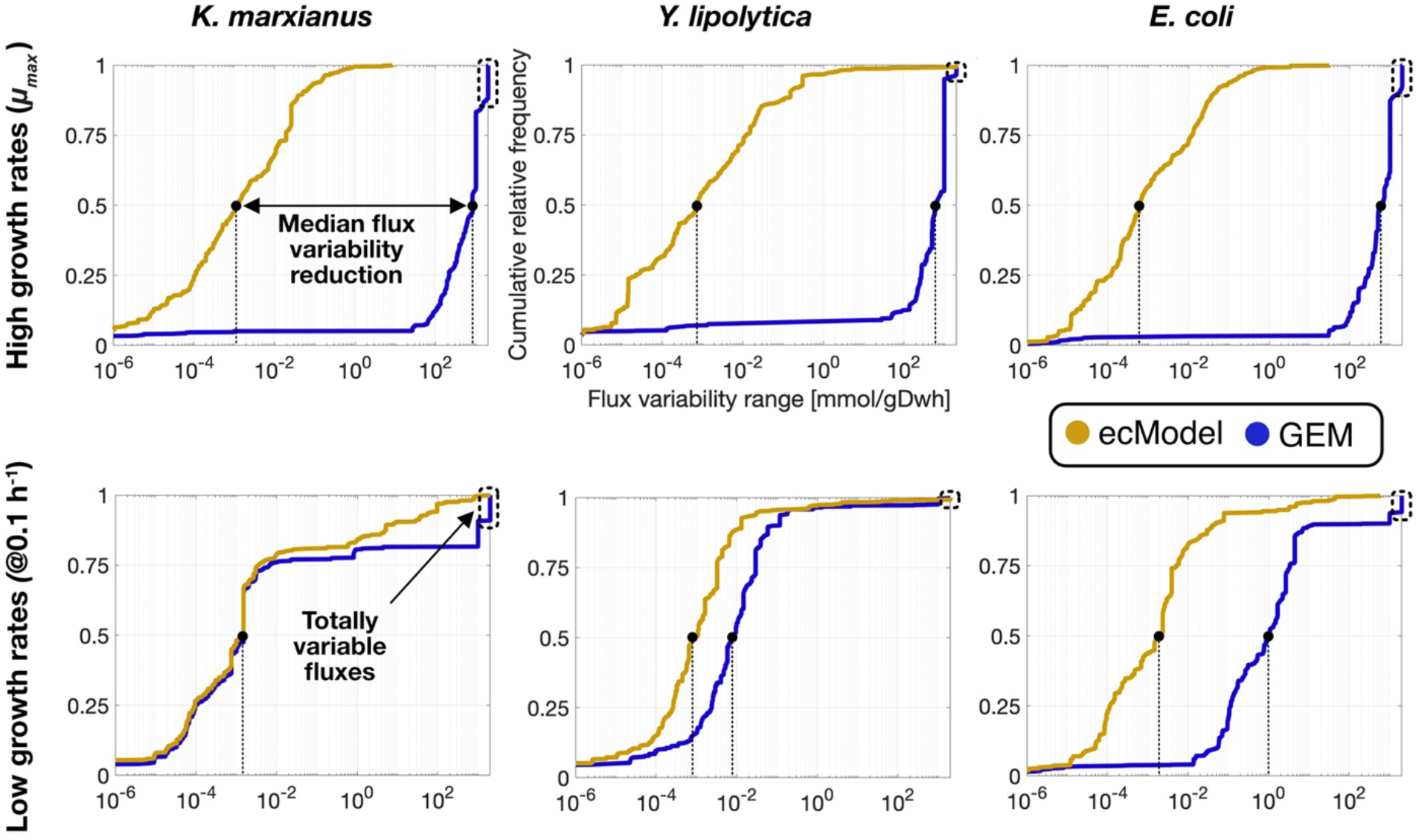
Cumulative distributions of flux variability ranges for *i*SM996, *i*Yali4 and *i*ML1515 compared to their respective enzyme-constrained versions at low and high growth rates.

Further analysis of the FVA results revealed that a reduction of at least 95% of the variability range was achieved for more than 90% of all active fluxes at high growth rates in all ecModel. Interestingly, the aforementioned flux variability metrics were overall improved even for the chemostat conditions, despite a higher degree of constraining (fixed low growth rate and optimal uptake rate), which restrains these models to an energy efficient respiratory mode (**Supp. file 4**).

### Discussion

Here we demonstrated how enzyme constrained models for diverse species significantly improve simulation performance compared to traditional GEMs. Furthermore, to enable the community to easily adapt this modelling approach, we upgraded the GECKO toolbox for enhancement of genome-scale models with enzyme and omics constraints to its version 2.0. Major improvements on the *k*_cat_ matching algorithm were incorporated into the toolbox, based on phylogenetic distance between the modeled organism and the host organisms for data queries, and an automated curation of *k*_cat_ numbers for over-constrained models were incorporated into the toolbox. Major refactoring of the GECKO toolbox enabled a generalization of the method, allowing the creation of high-quality ecModels for any provided functional GEM with minimal need for case-specific introduction of new code. Additionally, several utility functions were integrated into the toolbox in order to enable basic simulation purposes, accessible retrieval of enzyme parameters, integration of proteomics data as constraints, flux variability analysis and prediction of gene targets for enhanced production of metabolites. Overall, it was shown that these enhancements to the GECKO toolbox improve the incorporation of kinetic parameters into a metabolic model, yielding ecModels with biologically meaningful kinetic profiles without compromising accuracy on phenotype predictions.

Two major limitations of the first version of the GECKO toolbox were its specific customization to the *S. cerevisiae* model, Yeast7, and the need of extensive manual curation for generating an ecModel suited for FBA simulations; thus, its applicability to other GEMs was not a straightforward procedure. To overcome these limitations, we generalized the code with the aim of making GECKO a model-agnostic tool. The development of a procedure for automatic curation of kinetic parameters enabled the generation of functional ecModels with minimal requirements for experimental data. Recently, ecModels for 11 human cancer cell-lines were generated with this automated procedure, using Human1 as a model input and RNAseq datasets together with the tINIT algorithm^10^ to generate cell-line specific networks^2^. These ecModels were used for the prediction of cellular growth and metabolite exchange rates at different levels of added constraints, resulting in remarkable improvements in accuracy when compared with predictions of their original GEMs. This highlights one of the main advantages of ecModels: their capability of yielding biologically meaningful phenotype predictions without an excessive dependency on exchange fluxes as constraints.

In order to further showcase the functionality of the GECKO toolbox 2.0, a family of new high-quality ecModels were generated for *E. coli, Y. lipolytica, K. marxianus* and *H. sapiens,* based on the original GEMs *i*ML1515, *i*Yali4, *i*SM996 and Human1, respectively. Furthermore, we generated a self-hosted pipeline for continuous and automated generation and update of ecModels, ***ecModels container***, so that each of the currently available ecModels (ecYeastGEM, ec*i*ML1515, ec*i*Yali, ec*i*SM996 and ecHuman1) are integrated to it, providing a version-controlled and continuously updated repository for high-quality ecModels.

Absolute proteomics measurements for the budding yeasts *S. cerevisiae*, *K. marxianus* and *Y. lipolytica* grown under multiple environmental conditions, were incorporated as constraints into their ecModels by using the proteomics integration module added to the GECKO toolbox. Analysis of metabolic flux distributions revealed that net reaction fluxes predicted by GEMs are not significantly affected by the incorporation of kinetic and proteomics constraints, however the explicit integration of enzymes into ecModels extends the range of predictions of classical FBA and enables computation of enzyme demands at the reaction and pathway levels. It was found that incorporation of proteomics constraints does not affect enzyme demand predictions significantly for most of the active enzymes at low dilution rates across the simulated conditions. However, we observed that a diversified utilization of isoenzymes, enforced by proteomics constraints, increases the predicted total protein mass allocated to central carbon and energy metabolism, in comparison to optimal enzyme allocation profiles. This result suggests the relevance of metabolic robustness in contrast to optimal protein utilization for microbial growth under environmental stress and nutrient-limited conditions.

Incorporation of proteomics data allows the use of ecModels as scaffolds for systems-level studies of metabolism, providing a tool for uncovering metabolic readjustments induced by genetic and environmental perturbations, which might be difficult to elucidate by purely data-driven approaches, specially at conditions of relatively low changes at the transcript^69^ and protein levels^64^. For all studied stress conditions in this study, we identified upregulated proteins (increased abundance) that are needed to operate at high saturation levels in stress conditions, while showing low usage at reference conditions, creating lists of potential gene amplification targets for enhancing stress tolerance in three industrially relevant yeast species (**Supp. file 3**). Upregulation and high saturation of enzymes in amino acid and folate metabolism were found to be common across the studied organisms and stress conditions (**Fig. S3 D** and **Supp. file 3**). These results suggests that yeast cells display enzyme expression profiles that provide them with metabolic robustness for microbial growth under stress and nutrient-limited conditions, in contrast to an optimal protein allocation strategy that prioritizes expression of the most efficient and non-redundant enzymes.

Our results on drastic reduction of median flux variability ranges and the number of totally unbounded fluxes for ec*i*Yali, ec*i*SM996 and ec*i*ML1515, together with previous studies^1,2,38^, suggest that a major reduction of the solution space of metabolic models to a more biologically meaningful subspace is a general property of ecModels. However, flux variability is an intrinsic characteristic of metabolism; therefore, metabolic models with highly constrained solution spaces may exclude some biological capabilities of organisms, which are not compatible with the set of constraints used for the analysis (exchange fluxes, growth rates and even profiles of kinetic parameters, considered as condition-independent in ecModels).

Here, the predictive capabilities of ec*i*ML1515 and *i*JL1678 ME-model (both for *E. coli*) for cellular growth and global protein demands on diverse environments were compared. The major improvement in predicted maximum growth rates, together with a comparable performance on quantification of protein demands, shown by ec*i*ML1515 suggest that, despite its mathematical and conceptual simplicity, the GECKO formalism is a suitable framework for quantitative probing of metabolic capabilities, compatible with the widely used FBA method and without the need of excessive complexity or computational power. Nevertheless, ME-models provide a much wider range of predictions that explore additional processes in cell physiology with great detail. Direct comparison between the predictions of these modelling formalisms, suggest that ME-models performance can be improved by incorporation of either curated or systematically retrieved kinetic parameters that are suitable for the modelled organisms.

Simpler modelling frameworks that account for protein or enzyme constraints in metabolism, such as flux balance analysis with molecular crowding (FBAwMC)^16,17^, metabolic modelling with enzyme kinetics (MOMENT)^23^ and constrained allocation flux balance analysis (CAFBA)^21^, have also been developed and used to explore microbial cellular growth^16,17,21^ and overflow metabolism^16,23^. These methods have overcome the lack of reported parameters for some specific reactions either by incorporation of proteomics measurements and prior flux distributions^23^, manual curation and sampling procedures^16,17^ or even by lumping protein demands by functionally related proteome groups. In contrast, the new version of the GECKO toolbox provides a systematic and robust parameterization procedure, leveraging the vastly accumulated knowledge of biochemistry research stored in public databases, ensuring the incorporation of biologically meaningful kinetic parameters even for poorly studied reactions and organisms.

The applicability of these other simple modelling formalisms to models for diverse species is limited as none of these methods has been provided as part of a generalized model-agnostic software implementation. Recently, a simplified variant of the MOMENT method (sMOMENT) was developed and embedded into an automated pipeline for generation and calibration of enzyme-constrained models of metabolism (AutoPACMEN)^70^. The pipeline was tested on the generation of an enzyme-constrained version of the *i*JO1366 metabolic reconstruction for *E. coli*, which also showed consistency with experimental data. This work represented a step forward in the field of constrain-based metabolic modelling, as it contributed to standardization of model generation and facilitating their utilization and applicability to other cases.

However, due to the intrinsic trade-off between model simplicity and descriptive representation, a limitation of the sMOMENT method is its simplification of redundancies in metabolism, which just accounts for the optimal way of catalysing a given biochemical reaction, discarding the representation of alternative isoforms that might be relevant under certain conditions. In GECKO ecModels, all enzymes for which a gene-E.C. number relationship exists are included in the model structure. As traditional FBA simulations rely on optimality principles one could, in principle, expect the same predicted flux distributions by sMOMENT and GECKO ecModels. Nonetheless, the explicit incorporation of all enzymes in a metabolic network enables explanation of protein expression profiles that deviate from optimality in order to gain robustness to changes in the environment, as it has been recently shown by the integration of a regulatory nutrient-signalling Boolean network together with an ecModel for *S. cerevisiae*’s central carbon metabolism^71^.

In conclusion, GECKO2.0 together with the development of the automated pipeline ***ecModels container*** facilitates the generation, standardization, utilization, exchange and community development of ecModels through a transparent version-controlled environment. This tool provides a dynamic, and potentially increasing, catalogue of updated ecModels trying to close the gap between model developers and final users and reduce the time-consuming tasks of model maintenance. We are confident that this will enable wide use of ecModels in basic science for obtaining novel insight into the function of metabolism as well as in synthetic biology and metabolic engineering for design of strains with improved functionalities, e.g., for high-level production of valuable chemicals.

## Material and Methods

### Automation pipeline and version-controlled hosting of the ecModels container

The ecModels repository is used to version-control the pipeline code and the resulting models. The pipeline is restricted to 2 short Python files, whose role is to decide when models need to be updated based on a configuration file **config.ini**, and to consequently invoke the use of GECKO for each model. Updates are deemed necessary when either the underlying dependencies (i.e., GECKO, RAVEN and COBRA toolboxes, the Gurobi solver, and libSMBL) or the source GEMs are independently updated to a new version (release) in their respective repositories.

The pipeline is designed be automatic and to not require supervision. It was developed to work with both version-controlled GEMs and GEMs downloadable from a URL, updating the version in the configuration after a new ecModel is obtained. For easy review, the pipeline log is publicly available under the *Actions* tab of the GitHub repository. The computation is performed through a self-hosted GitHub runner, further leveraging the transparent nature of the GitHub platform and the *git* version-control system. The resulting ecModel and updated configuration are committed to the repository, with the changes being made available for review through a pull request. Additionally, the GECKO output is also replicated in the pull request body. The ***ecModels container*** thus continues the transparency and reproducibility of the source models.

### Quantification of absolute protein concentrations for *S. cerevisiae, Y. lipolytica* and *K. marxianus*

Total protein extraction for the strains *Saccharomyces cerevisiae* CEN.PK113-7D (standard, low pH, high temperature, osmotic stress), *Kluyveromyces marxianus* CBS6556 (standard, low pH, high temperature, osmotic stress) and *Yarrowia lipolytica* W29 (standard, low pH, high temperature) was conducted as described previously (**Supp. file 2**). Three reference samples (hereafter, ‘bulk’ samples), one per strain, were constructed by pooling 5 μg of each experimental sample. Aliquots of 15 μg of total protein extract from each sample (3 strains x 4 conditions x 3 replicates) and the three bulks were separated on onedimensional SDS-PAGE short-migration gels (11×1 cm lanes, Invitrogen, NP321BOX). Yeast proteins digestion was performed on excised bands from gel gradient and digested peptides of UPS2 (Sigma) were used as external standards for absolute protein quantification (more details in **Supp. file 2**). Four μl of the different peptide mixtures (800 ng for yeast peptides and 949 ng for bulks) were analyzed using an Orbitrap Fusion™ Lumos™ Tribrid™ mass spectrometer (Thermo Fisher Scientific).

Protein identification was performed using the open-source search engine X!Tandem pipeline 3.4.4^72^. Data filtering was set to peptide E-value < 0.01 and protein log(E-value) < −3. Relative quantification of protein abundances was carried out using the Normalized Spectral Abundance Factor (NSAF)^73^ and the NSAF values obtained from UPS2 proteins in bulk samples were used to determine the suitable regression curves that allowed the conversion from relative protein abundance into absolute terms. MS data is available online on public databases via the PRIDE repository^74^ with the dataset identifier PXD012836.

### Simulation of condition-dependent flux distributions

Simulation of cellular phenotypes for conditions of environmental stress at low dilution rates with GEMs were performed by first setting bounds on measured glucose uptake and byproduct secretion rates according to experimental data from previous studies on chemostats^64^. Then the biomass production rate was constrained (both upper and lower bounds) with the experimental dilution rate (0.1 h^−1^). Maximization of the non-growth associated maintenance pseudo-reaction was set as an objective function for the parsimonious FBA problem as a representation of the additional energy demands for regulation of cellular growth at non-optimal conditions. The same procedure was followed for simulations with ecModels constrained by a total protein pool. For the case of ecModels with proteomics constraints, the same set of constraints was used but the objective function was set as minimization of the total usage of unmeasured proteins, assuming that the regulatory machinery for stress tolerance is represented by the condition-specific protein expression profile.

### Prediction of microbial batch growth rates

Batch cellular growth was simulated by allowing unconstrained uptake of all nutrients present in minimal mineral media, enabling a specific carbon source uptake reaction for each case while blocking the rest of the uptake reactions and allowing unconstrained secretion rates for all exchangeable metabolites. Maximization of the biomass production rate was used as an objective function for the resulting FBA problem. For prediction of total protein demands on unlimited nutrient conditions, media constraints were set as expressed above and experimental batch growth rate values were fixed as both lower and upper bounds for the biomass production pseudo-reaction. The total protein pool exchange pseudo-reaction was then unconstrained and set as an objective function to minimize, assuming that when exposed to unlimited availability of nutrients the total mass of protein available for catalyzing metabolic reactions becomes the limiting resource for cells. The **solveLP** function, available in the RAVEN toolbox, was used for solving all FBA problems in this study.

## Code availability

The source code of the updated **GECKO toolbox** is available at: https://github.com/SysBioChalmers/GECKO. The code for **ecModels container** and the whole catalogue of updated ecModel files can be accessed at: https://github.com/SysBioChalmers/ecModels. All custom scripts for simulations included in this study can be found at: https://github.com/SysBioChalmers/GECKO2_simulations. All of these repositories are public and open to collaborative continuous development.

## Acknowledgements

We are grateful to Feiran Li, Raphaël Ferreira, Jonathan Robinson and all the GECKO users that have provided feedback for improving our toolbox and extending its range of applications and to the CHASSY project consortium for having motivated and supported this work. This project has received funding from the European Union’s Horizon 2020 Framework Programme for Research and Innovation – Grant Agreements No 720824 and 686070. This work was also supported by the Knut and Alice Wallenberg Foundation and The Novo Nordisk Foundation - Grant no NNF10CC1016517.

---

**Figure S1.**
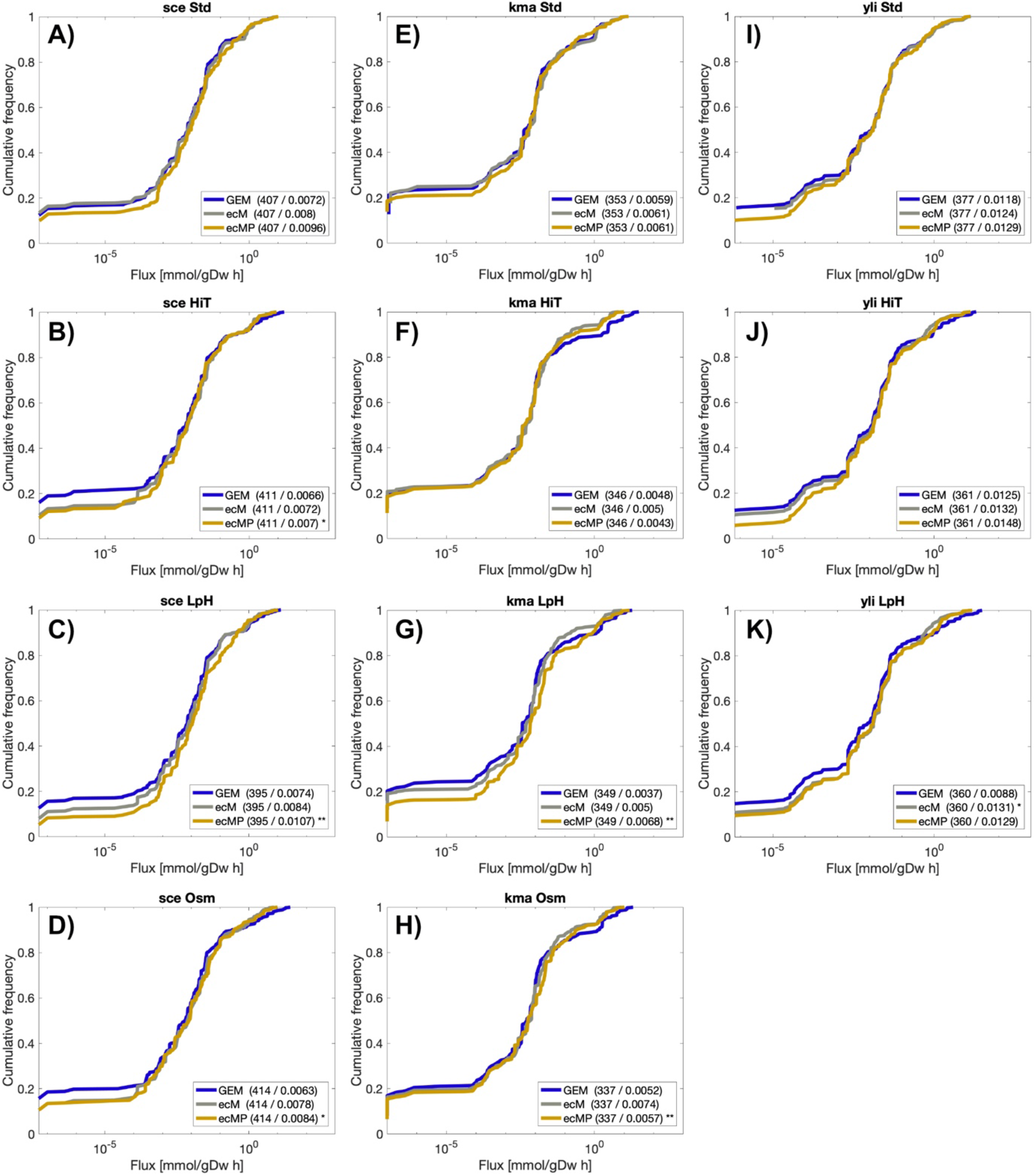
Cumulative distributions of metabolic fluxes. Flux distributions of ecModels were mapped to their corresponding reactions in the original GEMs and plotted together as cumulative distributions for all organisms and conditions. **A-D)** Cumulative distributions for *S. cerevisiae* models**; E-H)** Cumulative distributions for *K. marxianus* models; I-K) Cumulative distributions for *Y. lipolytica* models. Sample size and median flux values, in mmol/gDwh, are shown within parenthesis for all distributions in all the plots. Statistical significance under a two-sample Kolmogorov-Smirnov test between flux distributions for ecModels and their corresponding GEMs are shown as * (0.01 <= *p-value* < 0.05) and ** (*p-value<0.01).* sce – *S. cerevisiae*, kma – *K. marxianus*, yli – *Y. lipolytica*, std – Reference condition, HiT – High temperature condition, LpH – Low pH condition, Osm – Osmotic stress condition, GEM – Genome-scale metabolic model, ecM – ecModel with total protein pool constraint – ecP – ecModel with proteomics constraints.

**Figure S2.**
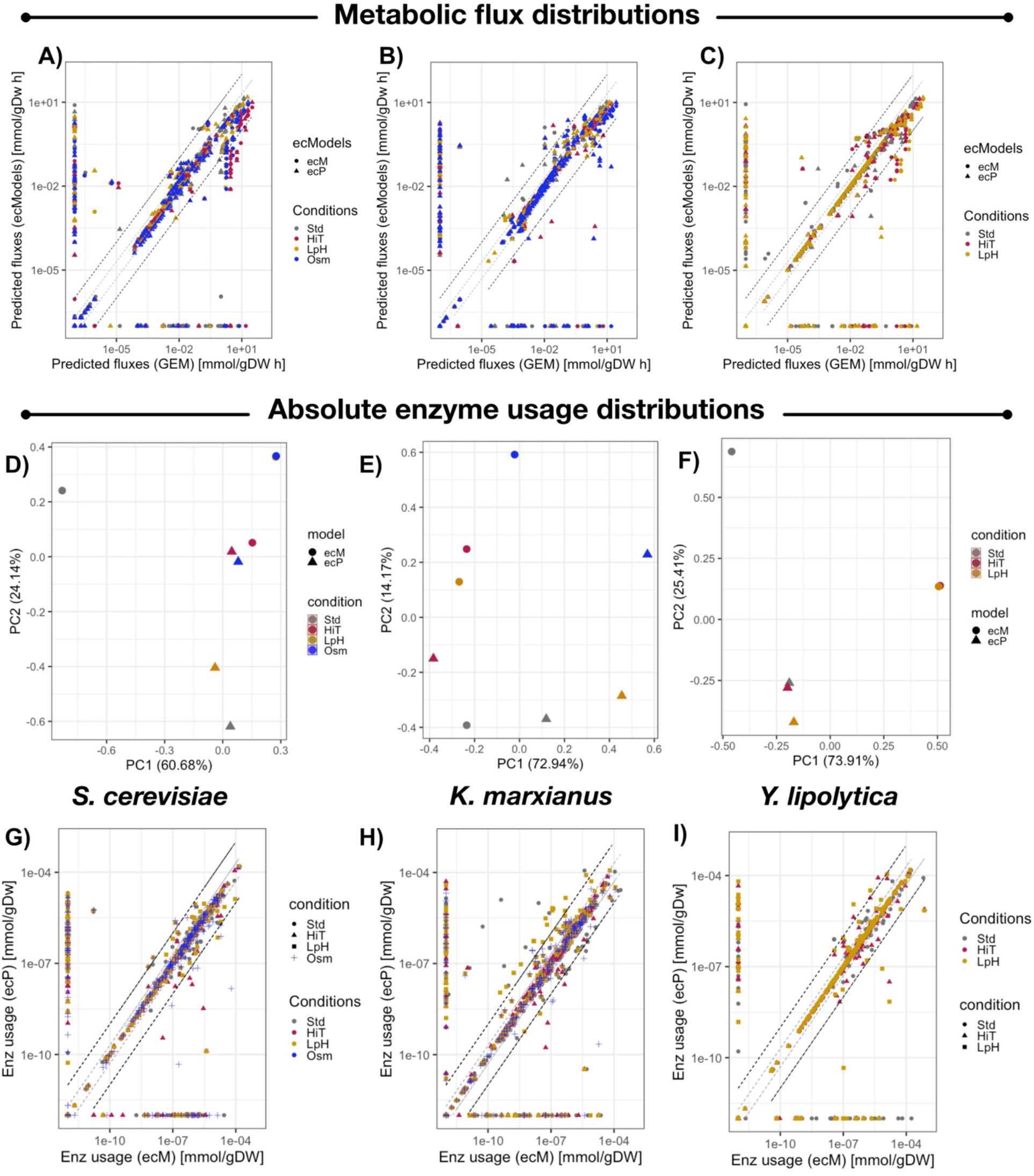
Comparison of predicted metabolic fluxes and enzyme usage distributions. Pairwise comparison of metabolic fluxes predicted by GEMs, ecModels and proteomics-constrained ecModels for **A)** *S. cerevisiae*, **B)** *K. marxianus* and **C)** *Y. lipolytica*. Principal component analysis on enzyme usage distributions predicted by ecModels and proteomics-constrained ecModels for **D)** *S. cerevisiae*, **E)** *K. marxianus* and **F)** *Y. lipolytica* subject to different environmental conditions. Pairwise comparison of enzyme usage profiles in mmol/gDw predicted by ecModels and ecModels with proteomics constraints for **G)** *S. cerevisiae* **H)** *K. marxianus* **I)** *Y. lipolytica*. Grey dashed lines indicate predictions in the interval 0.5 ≤ *fold change ≤* 2, whilst black dashed lines delimit the region of predictions within 0.1 ≤ *fold change ≤* 10, when comparing GEMs to ecModels (**A-C**) and ecModels to ecModels with proteomics data (**G-I**). Std – Reference condition, HiT – High temperatura condition, LpH – Low pH condition, Osm – Osmotic stress condition, GEM – Genome-scale metabolic model, ecM – ecModel with total protein pool constraint – ecP – ecModel with proteomics constraints.

**Figure S3.**
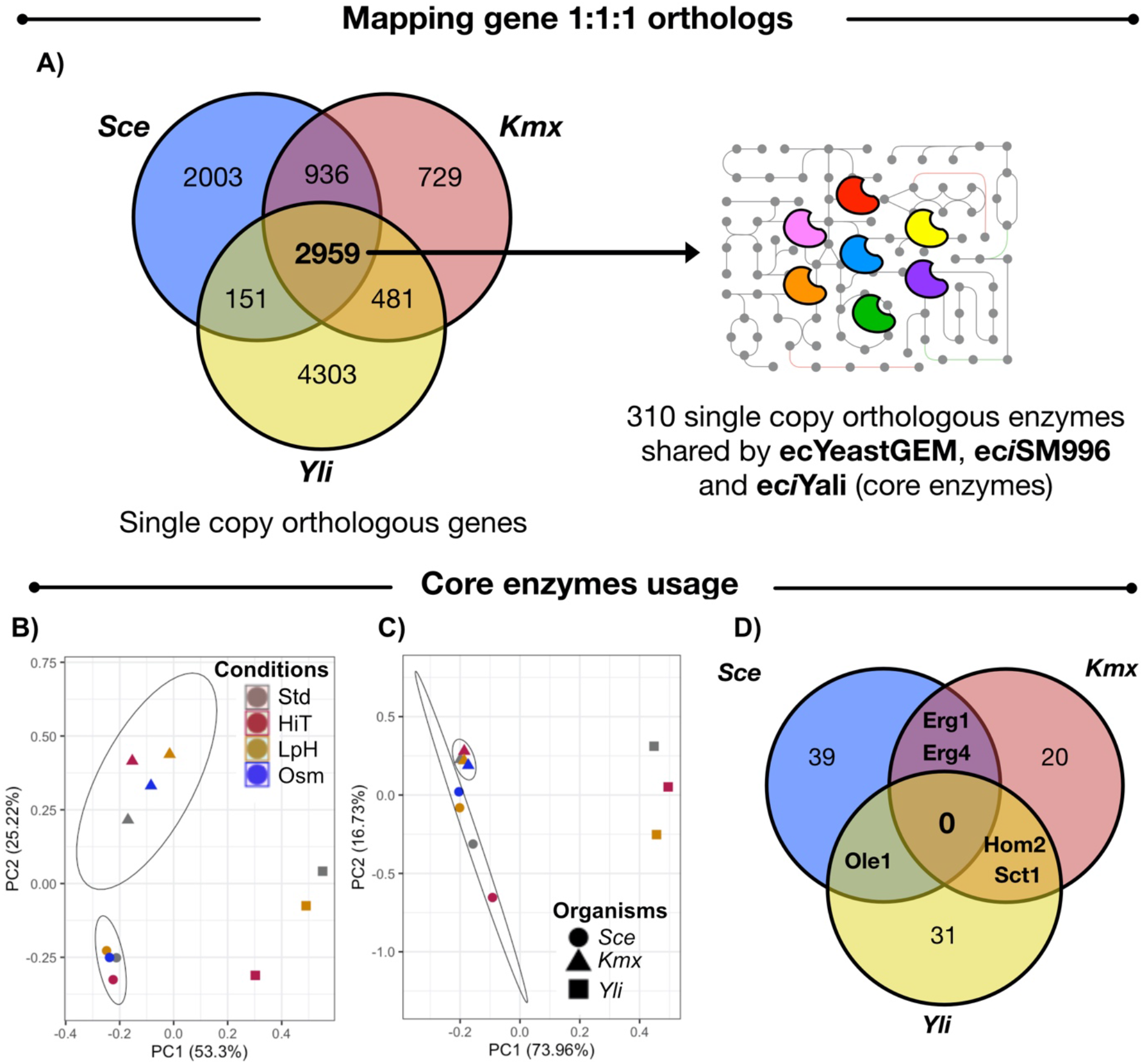
Evolutionary conserved enzymes across *S. cerevisiae, K. marxianus* and *Y. lipolytica.* **A)** Conservation of protein-coding genes amongst the three budding yeasts. Overlaps indicate number of genes conserved as single copy orthologs between yeast species. The uniprot codes for the 2,959 conserved protein-coding genes amongst the three species were mapped to their corresponding ecModels (ecYeastGEM, ec*i*SM996 and ec*i*Yali), 310 enzymes were found as single copy orthologs across the three ecModels. **B)** Principal component analysis on absolute abundances for the 310 core enzymes across the three yeast species for several experimental conditions. **C)** Principal component analysis on absolute enzyme usages predicted for the 310 core enzymes by ecYeastGEM, ec*i*SM996 and ec*i*Yali for several experimental conditions. **D)** Venn diagram for all core enzymes predicted as highly saturated (*relative usage* ≥ 0.95) in at least one environmental condition across the three yeast species. Std – Reference condition, HiT – High temperature condition, LpH – Low pH condition, Osm – Osmotic stress condition, Sce – *S. cerevisiae*, Kmx – *K. marxianus*, Yli – *Y. lipolytica.*

## Supplementary methods

### Improved *k*_cat_ matching algorithm

The *k*_cat_ matching algorithm in the GECKO toolbox queries kinetic parameters from BRENDA, the largest database available on enzymatic information^1^. However, one of the most important limitations to consider is that such parameters are only available for <10% of the known biochemical reactions^2^. The turnover number assignment to each of the enzymatic reactions present in a GEM is based on a flexible algorithm that allows the incorporation of kinetic parameters even when values for the specific organism and natural substrate of the enzyme are not available. However, as overestimation of microbial growth rates under environmental and genetic perturbations remains one of the main challenges for GEM development, biological relevance of the imposed kinetic constraints plays a crucial role for improving predictive accuracy^3^. In this regard, a global analysis for the reported *k*_cat_ values on BRENDA (**Supp. file 1**) pointed out the following potential issues.

1. The availability of kinetic parameters is highly heterogeneous, i.e. not all organisms have been studied to the same extent.
2. *k*_cat_ value distributions showed to be significantly different amongst kingdoms of life, therefore the catalytic activity of enzymes might be phylogenetically constrained.
3. *k*_cat_ value distributions are highly dependent on the metabolic context. For all kingdoms of life, there are important differences on the distributions for enzymes belonging to different metabolic pathways groups, being central carbon and energy metabolism enzymes the fastest group (on average) when compared to those involved in amino acid, fatty acid and nucleotide metabolism and secondary and intermediate metabolism.

In order to address the aforementioned limitations, the GECKO *k*_cat_ matching algorithm was modified aiming to provide a more accurate parameterization of models. A comparison between the introduced and previous hierarchical algorithms is shown in **Table S1.**

**Table S1:**
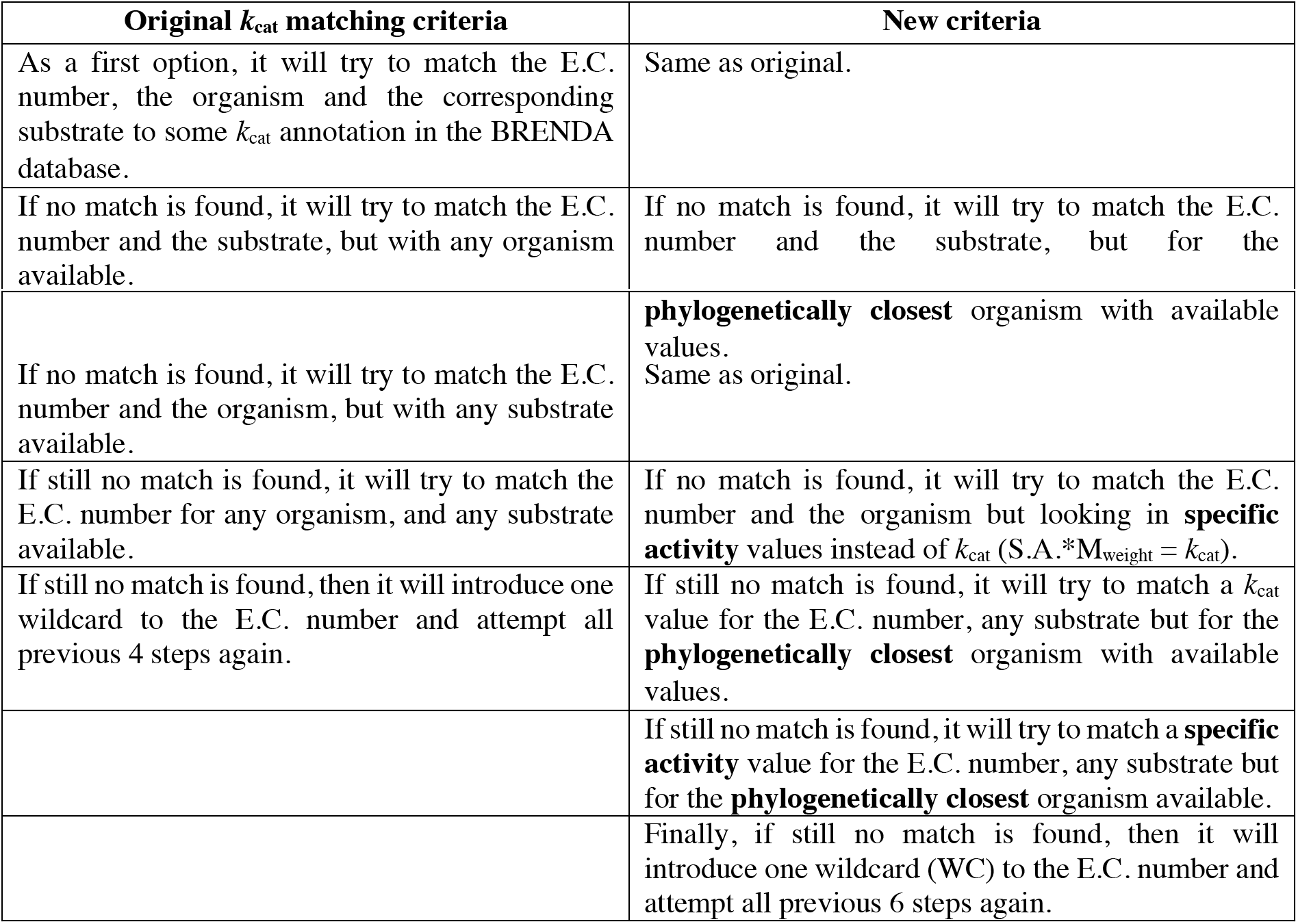
*k*_cat_ matching algorithms comparison.

### Estimation of phylogenetic distance between pairs of organisms

The phylogenetic distance between organisms is measured as the number of nodes of separation between two organisms in the KEGG taxonomical tree (incorporated as a MATLAB workspace file into the toolbox), this new feature follows from the assumption that kinetic parameters on enzymes have been finely tuned by evolution and are phylogenetically related^4^. The incorporation of specific activity values increases the parameter coverage and avoids the assignment of a high number of wild cards, making the assignments as close as possible to the original metabolic function of the specific enzyme class.

### Iterative curation of limiting *k*_cat_ numbers based on enzyme control coefficients

Once kinetic parameters and protein pool bounds have been incorporated into the ecModel it is very likely that overconstraining arises due to the intrinsic uncertainty of the incorporated *k*_cat_ values. For such cases, the module ***kcat_sensitivity_analysis*** flexibilizes the coefficients with a higher effect on the simulated objective function value based on enzyme control coefficients given by

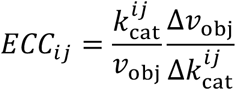

in which 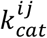 represents the *k*_cat_ parameter of the enzyme *i* in reaction *j*; *v_ob_*. is the original value in the objective function; 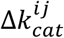 is an induced perturbation in the *k*_cat_ equivalent to 10-fold increase of its initial value; *Δv_obj_* is the change in the objective function after perturbing 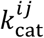.

The ECCs are ranked in a decreasing way and the enzyme with the coefficient is then selected for a 10-fold *k*_cat_ increase, based on the assumption that *k*_cat_ parameters may span orders of magnitude across organisms and substrates even for the same enzyme class. This procedure iterates until the ecModel is able to reach to provided experimental growth rate in the ***getModelParameters.m*** function. Information regarding the flexibilized *k*_cat_ values, their respective proteins and reactions, ECCs, flexibilized and original *k*_cat_ s is saved as a text file in the ***GECKO/model*** folder of the toolbox under the name ***kcat_modifications.txt***.

### Incorporation of proteomics constraints

The ***integrate_proteomics*** module in GECKO enables the generation of condition dependent models with proteomics constraints for any given dataset of absolute protein abundances [mmol/ gDw] with **m** replicates for **n** conditions. For each experimental condition, the abundance values are filtered, excluding proteins that are not present in at least 2/3 of the total number of condition replicates and also noisy measurements (proteins with relative standard deviation higher than 1 across replicates). Median values and standard deviations of abundance are calculated for each protein across replicates, then upper bounds are imposed on their corresponding enzyme usage pseudoreactions as follows:

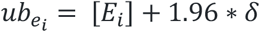

Where [*E_i_*] represents the abundance of the protein *i* in mmol/gDw and *δ* the standard deviation, the addition of 1.96 * *σ* accounts for a confidence interval of 0.95 in the protein abundance measurement. The experimental value for cellular growth rate at which the proteomics samples were obtained is then fixed as lower bound for the biomass pseudoreaction and measured fluxes on glucose uptake rate and, optionally, byproducts secretion rates are set as upper bounds for their respective exchange reactions (adding a numerical tolerance of 5%). The remaining total protein pool is then constrained by

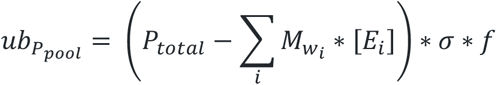

where *P_total_* is the measured total protein content in the cell in g_prot_/gDw; *M_wi_.* is the molecular weight of the measured protein *i; σ* represents an average saturation factor for the unmeasured enzymes, assumed as 0.5^5,6^; *f* accounts for the fraction that the unmeasured protein sector represents out of the total proteome in the cell, this value is calculated by using a paxDB proteome abundance file for the organism of interest as a reference, if no paxDB file is provided then a value of 0.5 is assumed.

Proteomics abundances are corrected for the oxidative phosphorylation complexes, trying to avoid overconstraining of potentially erroneously measured subunits that might limit the whole pathway. This correction is limited to just this pathway as it is desirable to modify the original dataset the least possible and abundance changes in OxPhos subunits are key to meet the phenotype energy requirements. Medium constraints are set by allowing free uptake of all compounds available in the culture medium and closing the rest of the uptake reactions. Additionally, all the upper bounds for production reactions (secretion of metabolites) are set to 1000 mmol/gDw h. Using an ***ecModel_batch*** with the same constraints setup, minimal enzyme requirements for the proteins present in the filtered dataset are retrieved from a parsimonious FBA solution vector. Enzyme abundances that are lower than the minimum requirements calculated by the FBA solution are corrected in the dataset.

A proteomics constrained ***ecModel_prot*** is obtained by the function ***constrainEnzymes.m***. If the model is overconstrained after imposing all the afore mentioned constraints, then the function ***flexibilizeProteins.m*** flexibilizes the top-limiting abundances (based on shadow prices for the measured proteins, given by: 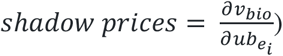 until the model is able to grow at the provided experimental growth rate. After this, an optimal enzyme usage profile compatible with the provided constraints is obtained and optimal levels are set as upper bounds for the flexibilized protein usages.

The total flexibilized mass of protein is drawn from the remaining protein pool (upper bound for protein_pool_exchange pseudoreaction) for consistency with mass conservation. Non-growth associated ATP maintenance is fitted according to condition specific experimental data if available (measurements on exchange fluxes of oxygen and CO_2_ from the same samples as the proteomics dataset). In the case of chemostat samples such conditions are set by first fixing the growth rate to the experimental value, minimizing the carbon source uptake, fixing its optimal value and then setting the total unmeasured enzymes usage as a new objective to minimize. Each condition-specific model is saved in ***GECKO/models/prot_constrained.***

### Comparative flux variability analysis

The function ***comparativeFVA.m*** in the FVA utitilities module provides a fair comparison of flux variability range distributions between a given GEM and its ecModel pair for glucose limited conditions (low dilution rates) and protein limiting regime (batch growth). For the chemostat case, a dilution rate of 0.1 h^−1^ was set as both lower and upper bound for the biomass pseudoreaction (+/− a tolerance value of 0.01%). Then the glucose uptake rate was set as an objective to minimize and its optimal value was then also fixed, using the same tolerance. Additional culture medium constraints were imposed (upper bound for exchange reactions of medium components were set to 1000 mmol/gDw h). All applied constraints were also applied to the original GEM. For the protein-limiting case, biomass production is maximized with the ecModel and then the optimal value is used to set a lower bound on the same reaction, in order to compare with the original GEM, the same optimal growth rate is fixed as both lower and upper bounds for the biomass pseudoreaction. A parsimonious flux distribution in which the total protein usage is minimized in the ecModel subject to all of the previous constraints is then obtained. For every reaction that is able to carry a non-zero flux in the original GEM (assessed by the RAVEN toolbox function ***haveFlux,m***) both minimization and maximization are performed for the original GEM. For the ecModel, such optimizations are performed on the governing pseudoreaction representing the same original reaction flux (i.e. arm reactions when isoenzymes are present), this is done for both the forward reaction and its reversible counterpart (if present). In order to avoid the introduction of artificial variability, the forward reaction is blocked when the backwards is optimized, and the same is applied to the opposite direction. For each reaction a flux variability range is given by

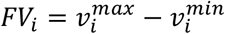

For the ecModel these ranges are given by

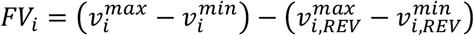

